# Organ-Specific Microbiomes of *Biomphalaria* Snails

**DOI:** 10.1101/2024.06.11.598555

**Authors:** Lauren V. Carruthers, Stephanie C. Nordmeyer, Timothy JC. Anderson, Frédéric D. Chevalier, Winka Le Clec’h

## Abstract

**Background:** The microbiome is increasingly recognized to shape many aspects of its host biology and is a key determinant of health and disease. The microbiome may influence transmission of pathogens by their vectors, such as mosquitoes or aquatic snails. We previously sequenced the V4 region of the bacterial 16S rRNA gene from the hemolymph (blood) of *Biomphalaria* spp. snails, vectors of the human blood fluke schistosome. We showed that snail hemolymph harbored an abundant and diverse microbiome. This microbiome is distinct from the water environment and can discriminate snail species and populations. As hemolymph bathes snail organs, we then investigated the heterogeneity of the microbiome in these organs.

**Results:** We dissected ten snails for each of two different species (*B. alexandrina* and *B. glabrata*) and collected their hemolymph and organs (ovotestis, hepatopancreas, gut, and stomach). We also ground in liquid nitrogen four whole snails of each species. We sampled the water in which the snails were living (environmental controls). Sequencing the 16S rRNA gene revealed organ-specific microbiomes. These microbiomes harbored a lower diversity than the hemolymph microbiome, and the whole-snail microbiome. The organ microbiomes tend to cluster by physiological function. In addition, we showed that the whole-snail microbiome is more similar to hemolymph microbiome.

**Conclusions:** These results are critical for future work on snail microbiomes and show the necessity of sampling individual organ microbiomes to provide a complete description of snail microbiomes.

## BACKGROUND

Over the past decade, microbiomes of both vertebrate and invertebrate hosts have been extensively characterized [1,2]. They are now recognized to shape many aspects of their host biology, health, and evolution [3,4]. As sequencing costs continue to decrease, investigating microbiomes at the organ or tissue level has become more affordable. While this approach is now routine for vertebrate microbiomes, microbiome studies of invertebrates are often limited to the whole body, particularly for smaller invertebrates (e.g., flies, mosquitoes, snails). However, using whole-body microbiome analysis can mask organ-specific and significant microbiome features due to its composite nature. Additionally, dynamic changes in microbiome composition can be obscured at the whole-body level. This issue shares similarities with challenges encountered in other fields of systems biology, such as cancer and immunology. For instance, the progress made from analyzing gene expression at bulk levels to single-cell levels has significantly transformed our understanding of biology and medicine [5,6]. To increase the resolution of microbiome studies, it is preferable to focus on organ or tissue-specific microbiomes rather than whole-body microbiomes. This approach has already been applied to invertebrate organisms, such as marine mollusks - oysters, mussels, and clams [7–9] - and terrestrial crustaceans [10,11].

*Biomphalaria* snails are freshwater mollusks and vectors of several trematode species, including the human parasite schistosome. Schistosomes infect over 200 million people in 78 countries in tropical and subtropical regions, with schistosomiasis responsible for over 200,000 estimated deaths annually [12–14]. Like many other marine and freshwater mollusks, *Biomphalaria* snails harbor a diverse microbiome in their hemolymph [15]. Since hemolymph bathes snail organs, our study aimed to investigate the composition and specificity of the microbiomes (i.e. bacterial communities) of these organs.

Therefore, we profiled the microbiome of multiple organs and tissues (hemolymph, stomach, gut, hepatopancreas (i.e. liver), and ovotestis) from two *Biomphalaria* species: *B. glabrata*, native to Brazil, and *B. alexandrina*, native to Egypt. We compared these specific organ microbiome profiles to whole-snail microbiomes, as well as the microbiomes found in the snail’s environmental water. Our findings demonstrate that the overall snail microbiome is not a true composite of the microbiomes of its various organs and is actually more similar to the hemolymph and, like the hemolymph, contains a highly diverse microbial community. Interestingly, we observed that the type of organ or tissue is the primary factor shaping the microbiome, rather than the snail species.

Our study highlights the importance of a well-thought-out sampling strategy when studying the snail microbiome. Both hemolymph and whole-snail samples capture a high microbiome diversity, but not in the same way. Conversely, the gut and stomach exhibited the highest microbiome density of all organs. Therefore, tailoring the sampling approach to the specific research question is crucial to provide the most accurate answer and avoid masking the effect of specific or rare taxa when analyzing the snail microbiome.

## METHODS

### 1. Biomphalaria spp. snail culture

*Biomphalaria alexandrina* and *B. glabrata* BS90 snails were reared separately in 10-gallon tanks containing aerated freshwater (sourced from a well on site at Texas Biomedical Research Institute) at 26-28°C on a 12-hour light – 12-hour dark photocycle. They were fed *ad libitum* with green leaf lettuce. *B. alexandrina* snails have been reared in the laboratory since 2012, originating from the Theodor Bilharz Research Institute in Egypt. *B. glabrata* BS90 snails were acquired from the Biomedical Research Institute Schistosomiasis Resource Center (Rockville, MD) and have been raised in our laboratory since 2013. All snails used in this study were adult specimens, with a shell diameter ranging between 10-15 mm.

For each species, we sampled fourteen snails from two independent tanks over two consecutive days. These snails were then transferred to clean plastic trays (20 x 6 x 7 cm) containing approximately 500 mL water taken from their respective tanks. They were starved overnight before organ and tissue collection. To mitigate the potential impact of the sampling time, we alternated between morning and afternoon whole-snail collection and dissections for each species (Fig. 1).

**Fig. 1.**
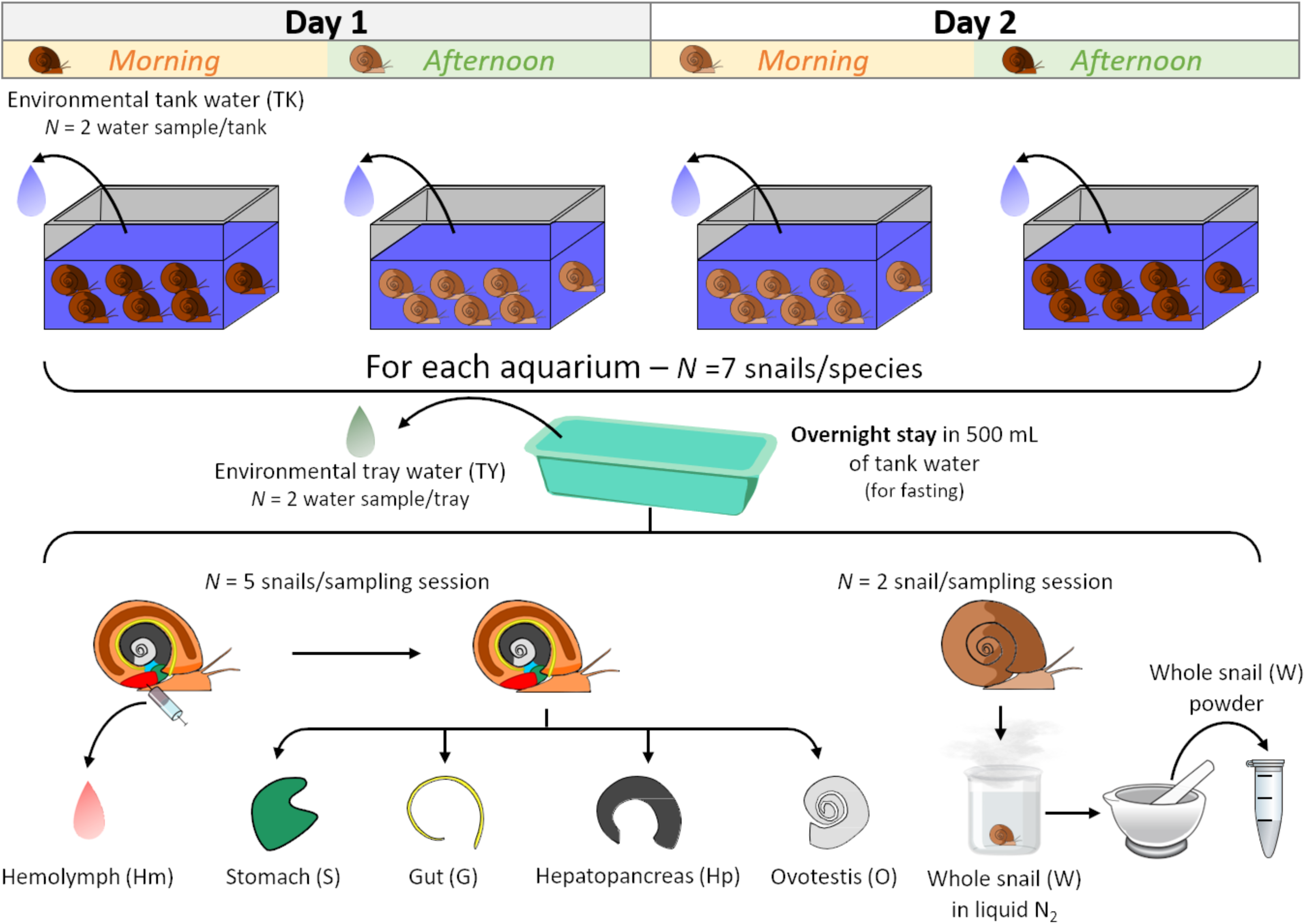
Experimental design. We sampled 14 *Biomphalaria alexandrina* (Ba) and 14 *B. glabrata* (BgBS90) snails from two different tanks alternately in the morning or the afternoon. We fasted the snails overnight in trays containing water from their respective tanks. For each batch, we sampled hemolymph, then stomach, gut, hepatopancreas (liver) and ovotestis from five snails, and snap-froze two whole snails. We sampled tank and tray waters as environmental controls. We extracted DNA from each sample to prepare 16S rRNA gene libraries, and sequenced these libraries on a MiSeq platform.

### 2. *Biomphalaria spp.* snail sample collection under sterile conditions

For each individual snail, we thoroughly disinfected the shell by wiping it three times with Kimwipes dampened with 70% ethanol [15]. After cleaning the shell, we collected hemolymph through cardiac puncture using a sterile 1 mL syringe and a 22-gauge needle. The collected hemolymph was immediately placed in a sterile 1.5 mL microtube pre-cooled on ice. We then gently crushed the snail shells using sterilized glass slides and removed shell fragments using sterilized forceps. From each snail (N = 10 snails per species) (Fig. 1), we isolated the ovotestis, hepatopancreas, gut, and stomach, using sterile tweezers and a binocular microscope. Once isolated, each organ was washed twice in sterile water in small sterile Petri dishes and transferred to sterile microcentrifuge tubes placed on ice. Samples were then snap-frozen in liquid nitrogen and stored at -80°C until processing.

To analyze the microbiome of whole snails, we snap-froze four intact snails (including their shells) from each species in liquid nitrogen. These frozen snails were individually crushed into fine powder using a pestle and mortar chilled with liquid nitrogen [16]. We transferred 100 µL aliquots of the powder to chilled microcentrifuge tubes before being stored at -80°C for gDNA extraction (Fig. 1).

Additionally, we collected two 500 µL aliquots of water from each tank where *Biomphalaria spp*. populations were raised, as well as from each tray in which the snails were housed overnight prior to dissection. These water samples were stored at -80°C and served as environmental controls.

### 3. DNA extraction of snail tissue

We extracted DNA from snail organs (ovotestis, hepatopancreas, stomach, or gut), whole-snail powder, and 40 µL of hemolymph samples, as well as 40 µL of water samples from tanks or trays, was extracted using the DNeasy Blood and Tissue kit (Qiagen). The extraction procedure followed the manufacturer’s protocol with the following modifications [15]. After addition of the ATL lysis buffer, we homogenized snail tissue samples (ovotestis, hepatopancreas, stomach, gut) using sterile micro-pestles. After addition of proteinase K, ovotestis, hepatopancreas, stomach, gut and whole-snail samples were incubated for 2 h in a water bath at 56°C, whilst hemolymph and water control samples were incubated for 1 h [15]. We eluted DNA in 50 µL of AE buffer for all the samples.

### 4. 16S rRNA gene library construction and sequencing

We prepared the 16S rRNA gene libraries in a laminar flow cabinet to avoid contamination. These were performed in triplicate to avoid potential biases introduced during PCR. We amplified ∼250 bp of the V4 region of the 16S rRNA gene from bacteria and archaea, using a set of specific primers (515f and 806rB) designed by the Earth Microbiome Project [17]. The 515f forward primers were barcoded to enable sample identification. Each reaction consisted of 4 µL of 1x 5 Prime Hot Master Mix (QuantaBio), 2 µL of sterile PCR grade H_2_O (MoBio Labs), 1 µL of 2 µM 515f barcoded primer, 1 µL of 2 µM 806rB primer (Supp. Table 1), and 2 µL of total gDNA template.

**Table 1.** ANOVA results of the linear mixed effect model on factors shaping microbiome diversity. We explored the impact of snail species and sample types (organs, hemolymph and whole snails) on three α-diversity indices. The most significant factor shaping the diversity is the sample type followed by the interaction between species and sample type and finally the species for Faith’s phylogenetic diversity and Simpson evenness.

|  |  | Observed richness |  | Faith's Phylogenetic diversity |  | Simpson evenness |  |
| --- | --- | --- | --- | --- | --- | --- | --- |
|  | Degrees of freedom | F value | p-value | F value | p-value | F value | p-value |
| Species | 1 | 2.98 | 0.09 | 4.68 | 0.04 | 0.22 | 0.64 |
| Sample type | 5 | 30.10 | 4.30e-17 | 25.99 | 1.59e-15 | 4.96 | 4.47e-04 |
| Species:Sample type | 5 | 1.61 | 0.18 | 1.52 | 0.19 | 5.12 | 3.37e-04 |

Libraries were amplified using the following thermocycler conditions: 95°C for 3 minutes, then 35 cycles of 95°C for 45 seconds, 50°C for 1 minute, 72°C for 90 seconds, then 72°C for 10 minutes. We electrophoresed 2 µL of each PCR product on agarose gel (2%) to check amplicon size, and to determine the absence of contamination in sterile water PCR controls. We pooled triplicate PCR products together (24 µL) in a 96-well plate and purified them by adding 44 µL of AMPure XP beads (Beckman Coulter). After 5-minute incubation, we placed the plates a plate magnetic device for 5 minutes. We then discarded the supernatant, and washed bead pellets twice in 200 µL of freshly-prepared 70% ethanol for 1 minute. After drying beads for 3 minutes, we removed the plate from the magnetic device, and added 20 µL of nuclease-free sterile water in each well for library elution. After a 2-minute incubation, the plate was placed on the magnetic device for 3 minutes. We then transferred each supernatant (i.e., purified barcoded libraries) into a fresh 96 well-plate. All incubations were done at room temperature. We quantified these libraries using the Picogreen quantification assay (Invitrogen) following the manufacturer’s instructions. Finally, we made an equimass pool of all the libraries using 15 ng of each library. We quantified the final pool fragment size using the 4200 TapeStation (DNA screen tape, Agilent Technology) and molarity using the Kappa qPCR library quantification kit for the Illumina sequencing platform (following the manufacturer’s recommendations). We sequenced the pool using an Illumina MiSeq platform (2 x 300 bp – v2 chemistry) at Texas Biomedical Research Institute (San Antonio, Texas). Raw sequencing data are accessible from the NCBI Sequence Read Archive under BioProject accession number PRJNA1113672.

### 5. Estimation of bacterial density by quantitative PCR

We quantified the number of bacterial 16S rRNA gene in each snail and environmental water sample using real time quantitative PCR (qPCR). While 16S rRNA gene copies can vary between and within bacterial taxa [18], we used this measure to estimate the bacterial density. The reactions were performed in duplicate using the QuantStudio 5 (Applied Biosystems) following the protocol outlined in [15]. We normalized the 16S qPCR data for hemolymph and environmental water samples as follows:

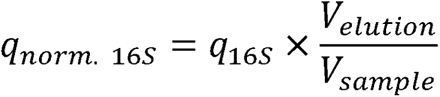

The bacterial density for whole-snail and organ microbiome samples may vary depending on the quantity of tissue sampled. Therefore, we normalized the 16S rRNA gene copy number to the number of snail cells. We estimated the number of snail cells by quantifying the single-copy gene *piwi* [19] in each sample and dividing this number by two as snails are diploid organisms. Real-time quantitative PCR for the *piwi* marker was conducted according to the methodology described in [16]. We normalized the 16S data as follows:

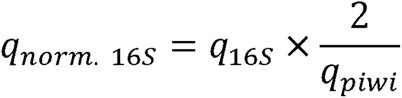

### 6. Data processing

Commands and scripts used for processing sequencing data and performing downstream analysis are available in a Jupyter notebook on Github (https://github.com/fdchevalier/Biomphalaria-organ-microbiomes).

We demultiplexed the raw sequencing data obtained from the MiSeq using bcl2fastq (v2.17.1.14, Illumina) with the default parameters. We used the fastq files generated as input for Qiime2 (v2019.10) [20]. We checked the read quality from the results of the demux module. We then denoised and clustered the sequences into amplicon sequence variants (ASVs) using the denoise-paired command from the dada2 module with a max expected error of 2. We determined the taxonomy of the ASVs using the SILVA database (release 138.2) and the classify-consensus-vsearch command of the feature-classifier module using a 97% identity threshold. ASVs with unassigned taxonomy were blasted against the NCBI nt database using megablast from blast+ (v2.9.0) to identify eukaryotic contaminants for removal in downstream analysis. We used a relatively lenient e-value threshold (1e-2) to increase our power of detection. To build a phylogenetic tree, we aligned the ASVs and masked the highly variable positions using the mafft and mask commands from the alignment module. Finally, we built and rooted a tree from the alignment using the fasttree and midpoint-root commands from the phylogeny module.

### 7. Statistical analysis

All statistical analyzes and graphs were performed using R software (v3.5.1). We used several packages to compute α-diversity statistics from the Qiime2 data: phyloseq [21], microbiome [22], and picante [23]. We compared α-diversity results using a pairwise Wilcoxon test. We used the following linear mixed-effect models to test which factor(s) influenced the α-diversity indices (1) and to test whether the collection period (morning/afternoon or day) had an impact (2):

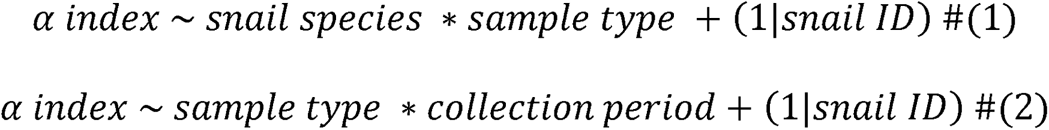

We compared model residues with ANOVA tests from the lmerTest package [24]. Water samples were excluded from the model.

We used the vegan [25] and the pairwiseAdonis [26] packages to perform β-dispersion and Permanova tests, respectively. We tested for significant differences using a *t*-test or a Wilcoxon test when data did not follow a normal distribution (Shapiro test, p < 0.05).

We compared bacterial density between sample types using a Kruskal-Wallis test (as the data did not follow a normal distribution; Shapiro test, p < 0.05), followed by pairwise Wilcoxon tests as post-hoc analysis. We used Kendall’s tau coefficient to test the correlation between the estimated 16S density in hemolymph and in organs (p-values < 0.05 were considered statistically significant).

## RESULTS

### 1. Experimental design and data summary

In the present study, we investigated the microbiomes associated with snail organs (gut, stomach, hepatopancreas, ovotestis) and compared them to the microbiomes of the hemolymph and whole-snail, as outlined in Fig. 1. We extracted DNA from each sample, and amplified and sequenced the V4 region of the 16S rRNA gene using an Illumina MiSeq platform. We obtained an average of 129,640 ± 3,782 reads per library (mean ± s.e.) (Supp. Table 2). After processing, we retained on average 61.61 ± 1.81% of the initial reads following sequencing error filtering, 61.01 ± 1.80% of the initial reads after denoising the data using the dada2 module, 56.48 ± 1.65% of the initial reads after merging reads and 54.45 ± 1.59% of the initial reads after removing chimeras. The sequencing depth was relatively even between sample types (Supp. Table 2). We performed rarefaction curve analysis to estimate how species richness changes with the number of reads subsampled from each library. Each curve reached a plateau, indicating that our sequencing effort effectively captured the microbiome diversity of the snail and their environment (Supp. Fig. 1).

Taxonomy assignment of the 16S rRNA gene sequences from all libraries revealed a total of 1,856 ASVs (amplicon sequencing variants). We excluded 22 (1.2%) ASVs that were assigned to mitochondria, chloroplasts or eukaryotes. Of the remaining 1,834 ASVs, 549 ASVs (127 without taxonomic assignment) were shared between *B. alexandrina* and *B. glabrata* BS90, 554 ASVs (of which 189 were without taxonomic assignment) were exclusively associated with *B. alexandrina,* and 556 (of which 219 were without taxonomic assignment) were exclusively associated with *B. glabrata*.

### 2. Snail organs exhibit their own microbiome

#### a) α**-diversity**

We detected a microbiome (i.e. the presence of bacterial communities) in each organ investigated (stomach, gut, hepatopancreas, ovotestis) as well as in hemolymph and whole snails. The α-diversity indices (within sample type diversity) showed similar trends across both snail species (Fig. 2, Supp. Fig. 2). Hemolymph and whole-snail harbored the highest number of observed ASVs, while the organs exhibited approximately half of this diversity. *B. glabrata* consistently showed similar microbiome richness in its organs compared to *B. alexandrina*, except in the hepatopancreas, which had lower diversity in *B. alexandrina* (Supp. Fig. 2). This observed richness covered a diverse range of phyla, as shown by the Faith’s phylogenetic diversity indices (Fig. 2). Hemolymph and whole-snail displayed the most phylogenetically diverse microbiomes, whereas the organ microbiomes were the least phylogenetically diverse, with the exception of the *B. glabrata* ovotestis. Despite this phylogenetic diversity, microbiome compositions were dominated by certain taxa, as indicated by the Simpson evenness index, which showed low taxa evenness within communities (Fig. 2). The evenness of *B. alexandrina* samples (i.e. organs/hemolymph/whole-snail) ranged widely, with the ovotestis displaying the most uneven microbiome, while the gut and hemolymph showed the least dominance. In contrast, the variation in evenness between *B. glabrata* samples was limited, with all samples being dominated by a few species.

**Fig. 2.**
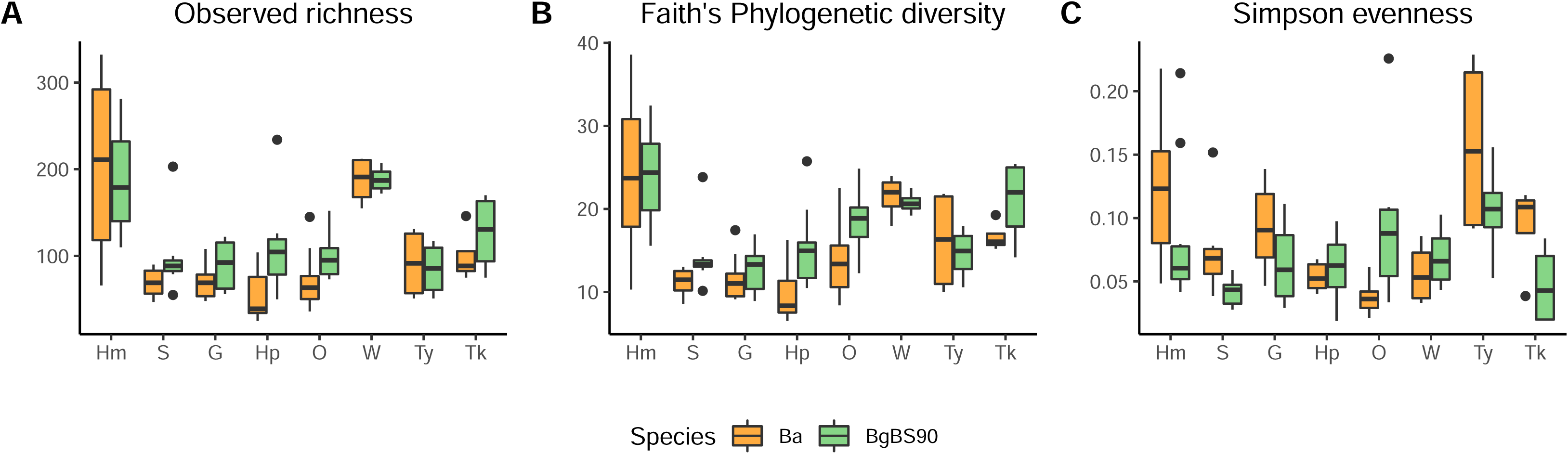
α-diversity indices of snail hemolymph, organs, whole snails, and water. We computed three α-diversity indices: observed ASVs (raw number of ASVs), Faith’s phylogenetic Diversity (based on phylogenetic distance between ASVs), and Simpson evenness (true evenness of taxa within a sample independently of taxa richness; a value of 0 means uneven taxa distribution in a sample). The highest diversity is found in hemolymph and whole snail samples. The two snail species (*B. alexandrina* (Ba) and *B. glabrata* (BgBS90)) showed differences in their diversity, especially between organs. All these bacterial communities are dominated by characteristic bacterial species. Hm: hemolymph, S: stomach, G: gut, Hp: hepatopancreas, O: ovotestis, W: whole snail, Ty: tray water, Tk: tank water. Each box plot represents the median and the interquartile ranges of the distribution. Statistical test is available in Supp. Table 3.

We used a linear mixed-effects model to better understand which factors (*snail species*, *sample type* [organs/hemolymph/whole-snail] or their interactions) predominantly shape the α-diversity indices (Table 1, Supp. Table 3). For both observed richness and Faith’s phylogenetic diversity indices, the *sample type* was the only factor with a significant effect (Table 1). For the Simpson evenness index, both *sample type* and the interaction between *sample type* and *species* had similar effects. This result was mainly driven by the hemolymph and whole-snails for observed richness and Faith’s phylogenetic diversity indices, and by the ovotestis for the Simpson evenness index (Supp. Table 3). However, it strongly suggests that the organ and tissue environments primarily shape the microbiome diversity, rather than snail species. We also tested the impact of *collection time* and *day* factors using a second model, which showed no significant effect (Supp. Table 3).

#### b) β-diversity

We compared the relatedness of the organ, hemolymph and whole-snail microbiomes, as well as environmental water microbiomes, using the UniFrac index (Fig. 3). This index measures the distance between two microbiome communities based on their composition and phylogenetic information [27]. The UniFrac index can be qualitative (presence/absence of taxa; unweighted UniFrac) or quantitative (including taxa abundance; weighted UniFrac). Differences in the unweighted UniFrac index indicate variations in the community composition associated with an environment, while differences in the weighted UniFrac index reflect variations in the community’s ability to thrive in that environment.

**Fig. 3.**
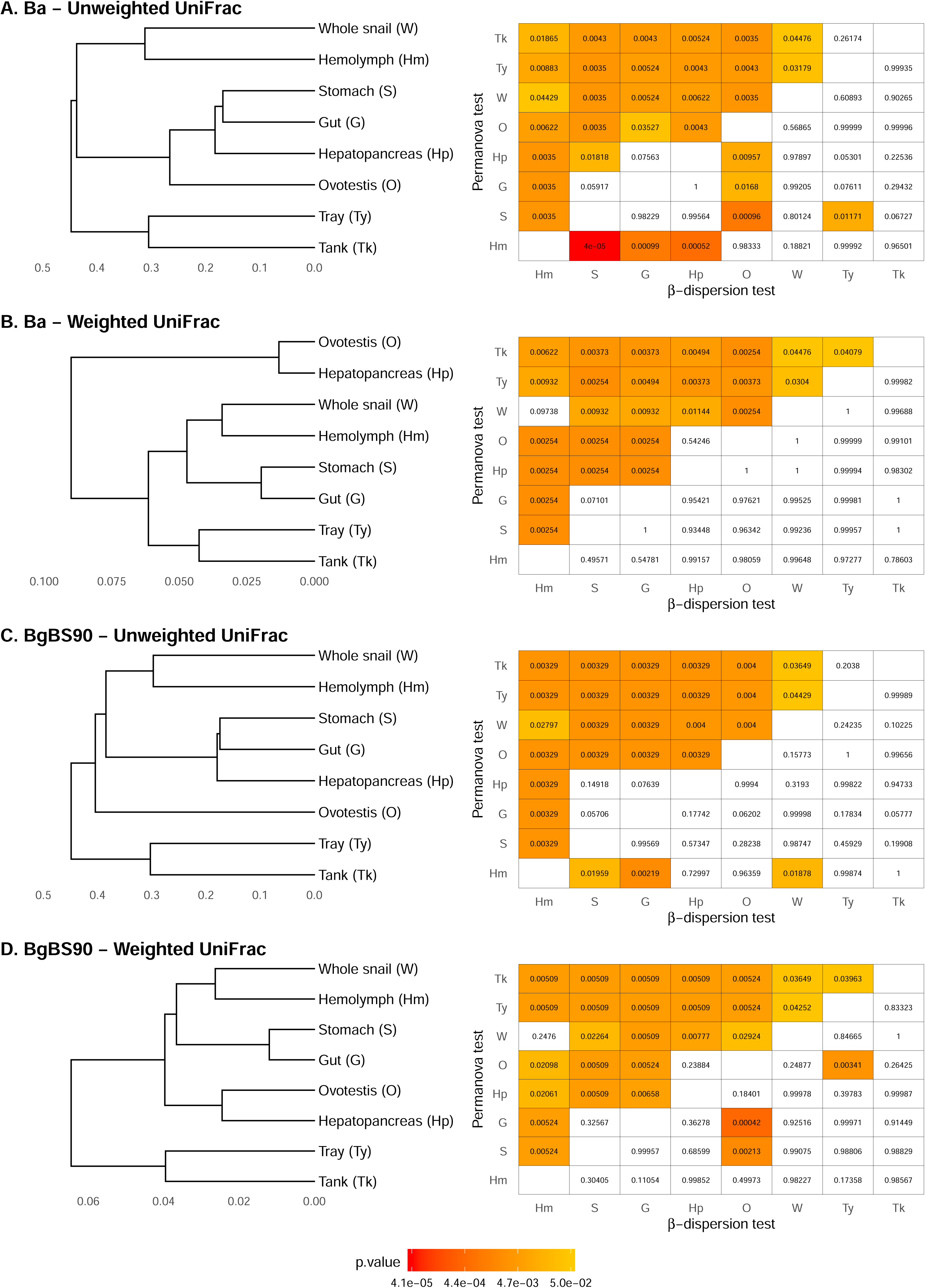
β-diversity indices of snail hemolymph, organs, whole snails, and water. We computed UniFrac indices: unweighted and weighted UniFrac. UniFrac measures account for the phylogenetic distance of the microbial taxa when measuring distances between groups. These measures can be weighted by the abundance of each microbial taxa. The matrices on the right side of each index showed the significance level of the Permanova test (differences between sample types) and the β dispersion test (homogeneity of variance between sample types confirming that significant PERMANOVA results are due to differences in distance between groups and not to differences in sample dispersion between groups). Organ microbiomes tend to cluster by functions. Whole-snail microbiomes are similar to hemolymph microbiomes. Ba: *B. alexandrina*, BgBS90: *B. glabrata*; Hm: hemolymph, S: stomach, G: gut, Hp: hepatopancreas, O: ovotestis, W: whole snail, Ty: tray water, Tk: tank water.

Overall, microbiome communities tended to cluster based on organ function and the physical distance between these organs [28], as demonstrated by both unweighted and weighted UniFrac measures in both snail species (Fig. 3). For instance, stomach and gut microbial communities consistently clustered together, suggesting shared similarities in both composition and abundance. The hepatopancreas, which performs a liver-like function, clustered with the digestive system in unweighted UniFrac analysis, suggesting the colonization of this organ by specific taxa from the digestive system. This may be due to the physical proximity of the gut and hepatopancreas. However, in weighted UniFrac analysis, the hepatopancreas consistently clustered with the ovotestis. Given the physical connection between the hepatopancreas and the ovotestis, this observation likely reflects the influence of the neighboring ovotestis microbiome, with some major taxa possibly colonizing the hepatopancreas. Interestingly, hemolymph and whole-snail consistently clustered together in both UniFrac measures, suggesting substantial overlap between taxa found in whole-snail and those in the hemolymph. However, the relative distance between hemolymph/whole-snail cluster and the organ clusters varied. In unweighted UniFrac analysis, this cluster was either farther (in *B. alexandrina*) or closer (in *B. glabrata*) to all other organs, suggesting the presence of specific taxa unique to this cluster. In weighted UniFrac analysis, the hemolymph/whole-snail cluster tended to be closer to the digestive system cluster, suggesting that both communities share abundant taxa. Environmental samples (tray and tank waters) formed a distinct cluster, consistent with previous observations [15].

We performed PERMANOVA tests to assess the robustness of the differences observed between sample types and clusters. Firstly, we confirmed that these differences were actually due to differences in group centroid distances rather than group dispersion by conducting a β-dispersion test (Fig. 3C). Groups showed similar dispersion for each comparison with a few notable exceptions: comparisons between hemolymph and the digestive system using unweighted UniFrac, and between the ovotestis and the digestive system using either unweighted UniFrac (*B. alexandrina*) or weighted UniFrac (*B. glabrata*), which showed significant differences. This suggests that the differences in PERMANOVA could be due to variation in group dispersion in addition to group distance, as indicated by the PCoA plots (Supp. Fig. 3). PERMANOVA tests revealed statistically significant differences in group distances for nearly all comparisons. This finding indicates that microbiomes are specific to their respective organs, determined by both the type of taxa associated (unweighted UniFrac) and the proportion of these taxa (weighted UniFrac). A few group distances showed consistent non-statistical differences: microbiomes of whole-snail and hemolymph, stomach and gut, and ovotestis and hepatopancreas were similar using weighted UniFrac for both snail species, while microbiomes of the stomach and gut, and the hepatopancreas and gut were similar using unweighted UniFrac for both snail species. These results suggest that these combinations share similar microbial communities. Distances between tank and tray water microbiomes revealed similar compositions (unweighted UniFrac) but different taxa abundances (weighted UniFrac).

#### c) Taxonomic diversity

As observed in the β-diversity analyses, hemolymph and organs exhibited distinct taxonomic microbiome compositions, with similarities either related to organ functions (such as stomach and gut microbiomes) or spatial localization (such as hepatopancreas and ovotestis) (Fig. 4). Additionally, hemolymph and whole-snail microbiomes showed strong similarities, which were consistent in both *B. alexandrina* and *B. glabrata*.

**Fig. 4.**
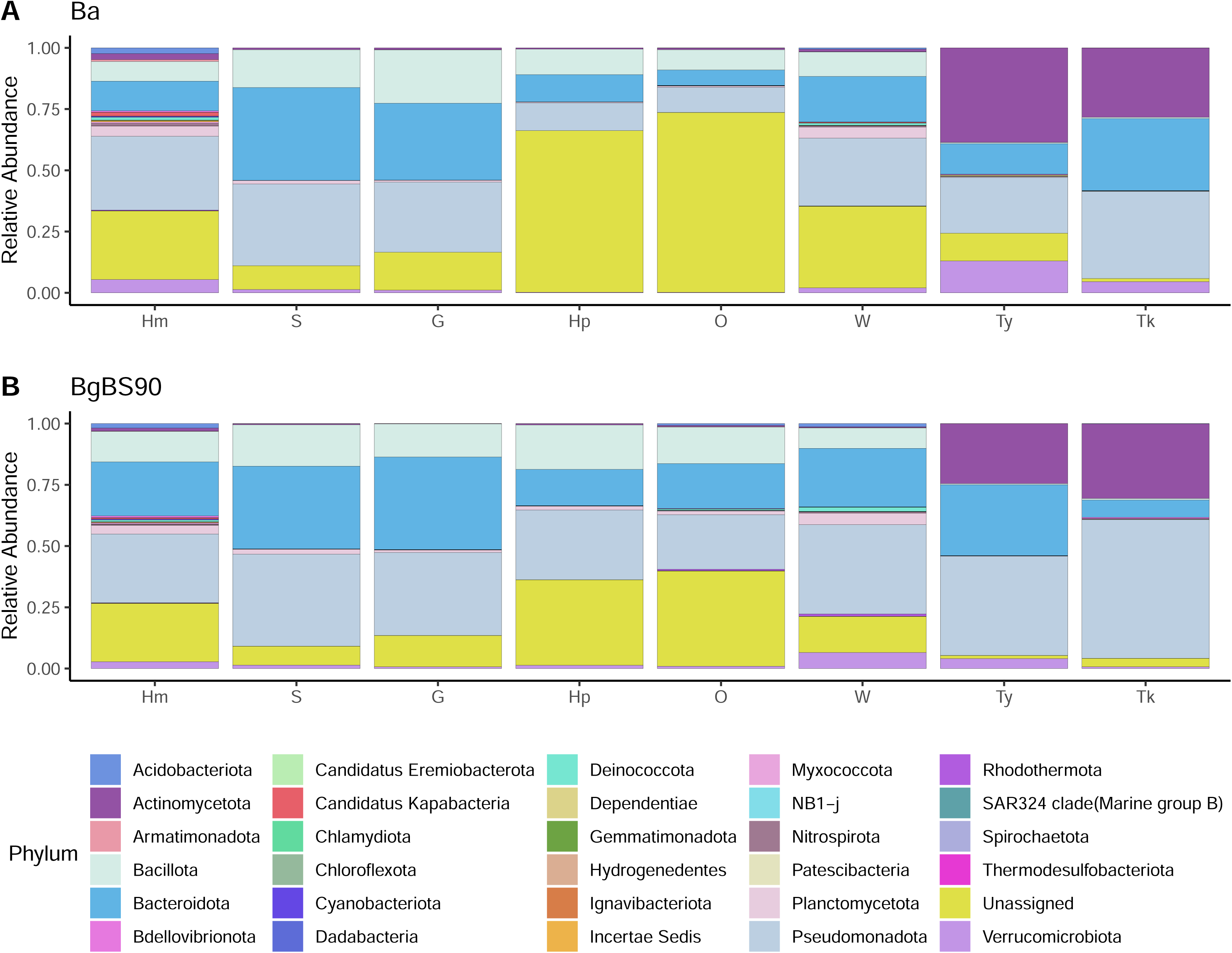
Taxonomic diversity of sample types in each snail species. Relative abundance of the different phyla in each sample type for each snail species (A & B). Composition varies between sample types with Bacteroidetes, Pseudomonadota (Proteobacteria), Tenericutes and unassigned phyla dominating snail sample types, and with Actinomycetota (Actinobacteria) and Pseudomonadota phyla dominating water samples. Composition is more similar between organs sharing functions (stomach and gut) or location (hepatopancreas and ovotestis). Proportion of unassigned phyla is high in snail sample types, particularly in *B. alexandrina* hepatopancreas and ovotestis. Ba: *B. alexandrina*, BgBS90: *B. glabrata*; Hm: hemolymph, S: stomach, G: gut, Hp: hepatopancreas, O: ovotestis, W: whole snail, Ty: tray water, Tk: tank water.

Hemolymph and whole-snail samples were mainly dominated by the phyla Pseudomonadota (formerly Proteobacteria) and Bacteroidetes. The two main classes of the Pseudomonadota phylum, Alphaproteobacteria and Gammaproteobacteria, were found in relatively similar proportions in our samples (Supp. Fig. 4). Our previous study on hemolymph showed dominance of Bacteroidetes in *B. alexandrina* [15], which was not observed here, suggesting that hemolymph microbiome composition is dynamic. The microbiomes of hemolymph and whole-snail varied mainly in the proportions of less-dominant phyla, including Acidobacteria, Actinomycetota (formerly Actinobacteria), Tenericutes, unassigned taxa and Verrucomicrobia. However, only Acidobacteria and Actinomycetota were consistently found in higher proportions in the hemolymph in both *B. alexandrina* and *B. glabrata*. Stomach and gut microbiomes were mainly dominated by Bacteroidetes, Pseudomonadota (mainly Gammaproteobacteria class) and Tenericutes phyla. Hepatopancreas and ovotestis microbiomes were strikingly dominated by bacteria from unassigned phyla, especially in *B. alexandrina,* where these phyla comprised over 50% of the community. These two organs may represent a very specific environment for poorly characterized bacteria. Finally, environmental water microbiomes were dominated by the phyla Actinomycetota, Bacteroidetes, and Pseudomonadota (predominantly Gammaproteobacteria class), similar to our previous observation [15], although Actinomycetota were found in lower proportions.

### 3. Shared microbes between hemolymph, organs and water and between snail species

Taxonomic diversity revealed shared microbiome community patterns between sample types. To explore this further, we investigated how many taxa were actually shared between the sample types (Fig. 5, Supp. Fig. 5). While sample types showed numerous distinct ASVs (especially in hemolymph and ovotestis), approximately 45% of the ASVs were shared between two or more sample types in each snail species. ASVs shared only between one snail sample type (organ or hemolymph) and whole snail may correspond to either (i) distinct ASVs from that sample type or (ii) shared ASVs detected in that sample type and another unsampled compartment (e.g., tegument) from the whole snail.

**Fig. 5.**
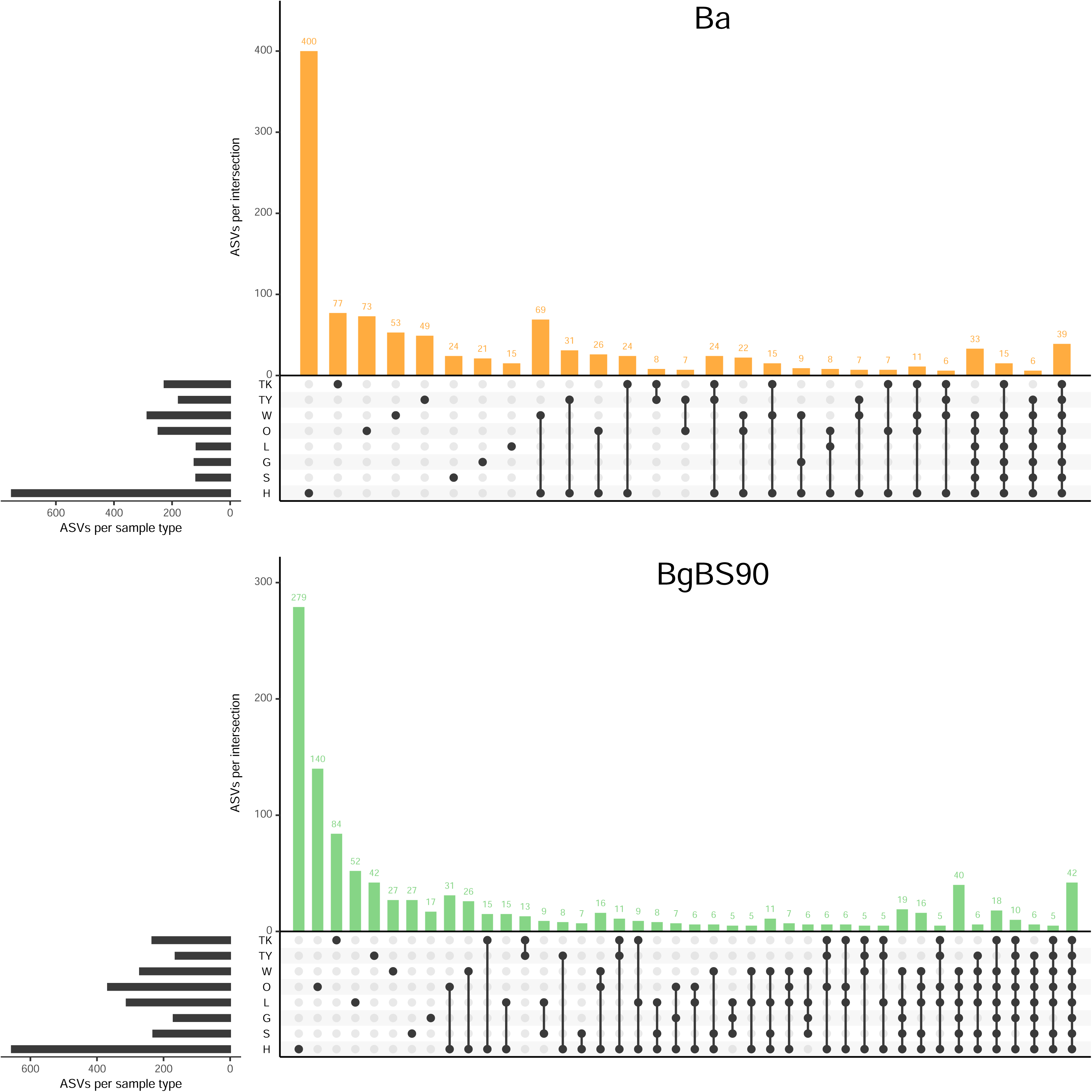
Shared ASVs between sample types by species. These upset plots show the number of ASVs (left horizontal barplot) and the number of ASV (upright barplot) for each intersection (dot plot) for *B. alexandrina* (Ba) and *B. glabrata* (BgBS90). For readability, we removed intersections with less than five ASVs. The full intersection set is available in Supp. Fig. 5. A significant proportion of ASVs are distinct, mainly found in the hemolymph and ovotestis. Some ASVs are also found ubiquitously in the snails but not in the environment. Hm: hemolymph, S: stomach, G: gut, Hp: hepatopancreas, O: ovotestis, W: whole snail, Ty: tray water, Tk: tank water.

A few trends were consistently observed between snail species. Among the shared ASVs, 40% to 50% were shared between hemolymph and ovotestis (Supp. Fig 6). Nearly half of the total shared ASVs (*B. alexandrina*: 47%, *B. glabrata*: 40%) were shared between hemolymph and water samples (trays or tanks) suggesting that water serves as a source or a recipient of these ASVs. In contrast, a similar comparison between the digestive system (gut or stomach) and water samples revealed that only about a quarter of the total shared ASVs were present in both compartments (*B. alexandrina*: 22%, *B. glabrata*: 26%), suggesting a relatively tight regulation of the microbiome in the digestive tract. A large proportion of shared ASVs (*B. alexandrina*: 67%, *B. glabrata*: 67%) were exclusive to snail samples (either whole-snails or organs / hemolymph) and not found in water samples. Around 13% of these ASVs were shared across all snail sample types, while about 7% of shared ASVs were ubiquitous, found in all snail and water sample types. Strikingly, stomach and gut microbiomes, which showed similar composition at the phyla level, shared only between 26% to 31% of ASVs (Supp. Fig 6). Hepatopancreas and ovotestis exhibited a similar trend (25% to 36%), despite being physically connected. This suggests that specific ASVs are adapted to particular organs. Approximately 70% of ASV intersections contained 4 or fewer ASVs (Supp. Fig. 5), which may be due to noise or detection limits in some compartments.

We also examined the number of ASVs shared between snail species for each sample type (Fig. 6). *B. alexandrina* and *B. glabrata* shared almost half of the ASVs found in their hemolymph or water samples. However, organs of *B. glabrata* had 1.5 to 5 times more ASVs than those of *B. alexandrina*, which aligns with the greater observed richness in *B. glabrata* compared to *B. alexandrina* (Fig. 2). Whole-snail samples showed more shared ASVs than distinct ones. These distinct ASVs may correspond to unsampled sections of the snails, such as the tegument, which is present in whole-snail samples.

**Fig. 6.**
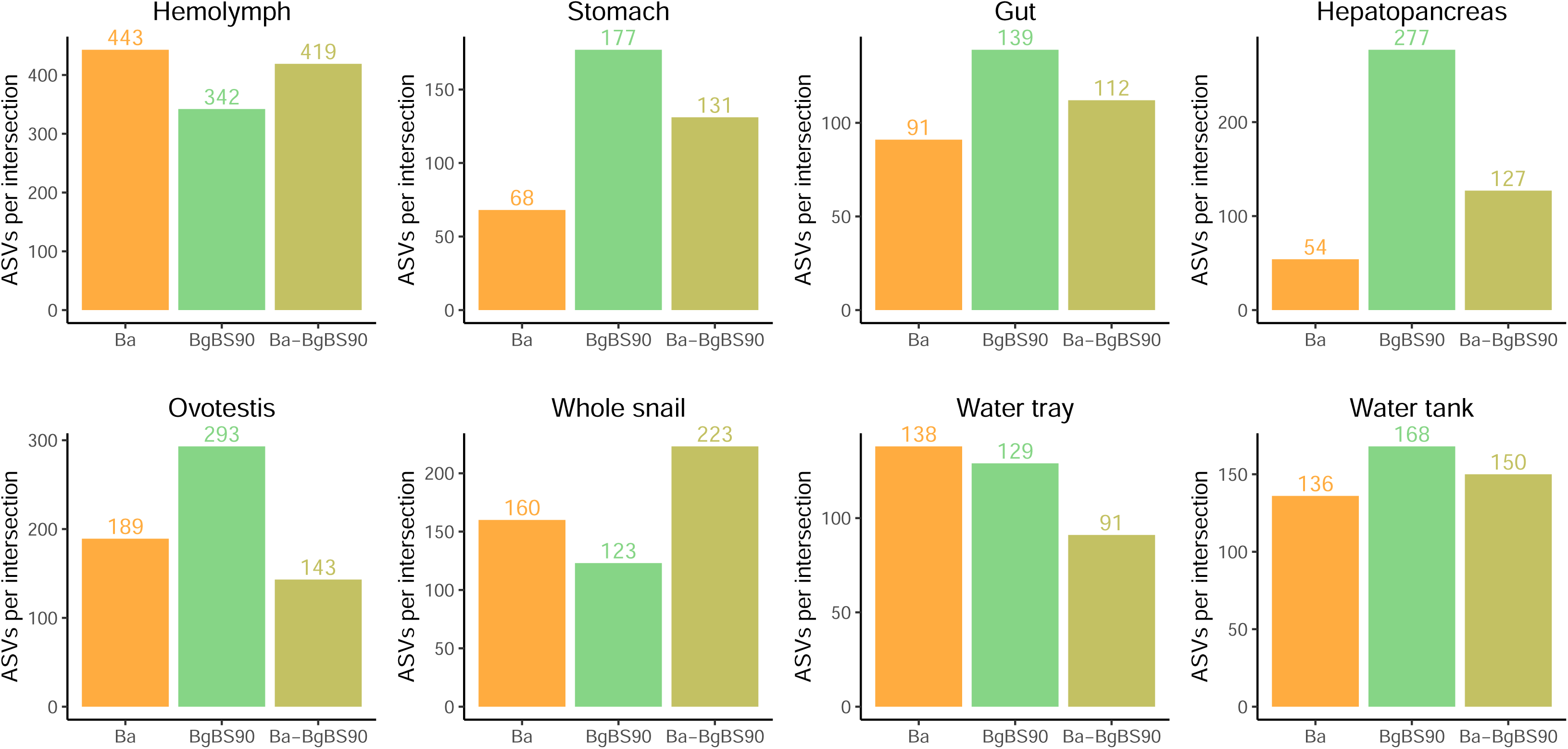
Shared ASVs between species by sample types. Barplots showed the number of distinct ASVs by species or shared between species for each sample type. Shared ASV between the two species are usually representing between a third to half of the ASV of a given sample type. *B. glabrata* (BgBS90) tends to have more distinct ASVs than *B. alexandrina* (Ba).

### 4. Snail bacterial density varies between organs

We estimated relative bacterial densities in hemolymph, organs and whole-snails by quantifying the 16S rRNA gene copy number using qPCR (Fig. 7, Supp. Fig. 7). Copy numbers were normalized either by volume extracted (hemolymph, water) or number of snail cells (organs and whole-snails). The bacterial densities observed in hemolymph (*B. alexandrina*: 3,227 ± 467 16S copies/µL (mean ± s.e), *B. glabrata*: 6,301 ± 1,009 16S copies/µL) were consistent with those previously quantified [15]. Four out of twenty hemolymph samples had densities higher than 10,000 copies/µL (*B. alexandrina*: 2/10 and *B. glabrata*: 2/10), with one significant outlier at around 263,500 copies/µL which was excluded from the analysis. The bacterial densities observed in organs and whole snails followed consistent patterns, except for the hepatopancreas. The stomach and gut microbiomes had the highest bacterial densities (*B. alexandrina*: 87.42 ± 7.31 16S copies/cell, *B. glabrata*: 136.89 ± 16.48 16S copies/cell), while the ovotestis had the lowest (*B. alexandrina*: 2.57 ± 0.30, *B. glabrata*: 0.78 ± 0.12 16S copies/cell). The bacterial densities in the hepatopancreas showed intermediate levels with strong differences between snail species. These differences could be biological or artefactual due to gut remnants remaining intertwined within the hepatopancreas. Whole-snails exhibited a limited number of 16S copies per cell, with a similar pattern between both species (*B. alexandrina*: 2.12 ± 0.77, *B. glabrata*: 4.21 ± 1.07 16S copies/cell). This overall low bacterial density is likely due to the head-foot region of the snail body, which constitutes a high proportion of the whole-snail sample but likely has low bacterial content.

**Fig. 7.**
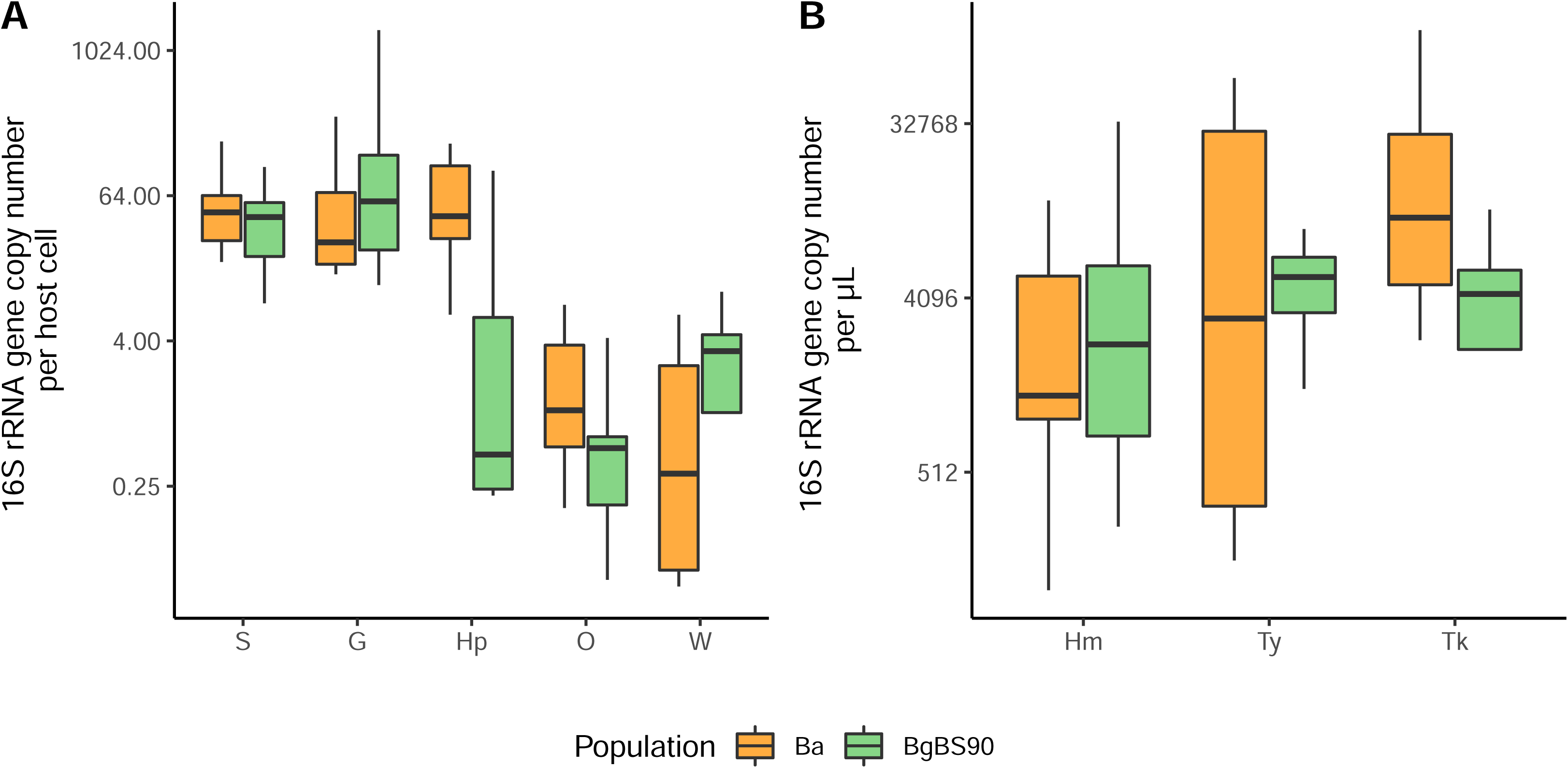
Comparison of 16S density between snail organs, whole snail, hemolymph and water. We quantified raw 16S copy number either host cell (organ and whole snail) or per microliter of hemolymph and water using qPCR. Stomach and gut had the highest densities, ovotestis the lowest, with species-specific variations in hepatopancreas densities. Hemolymph densities were consistent with previous studies. S: stomach, G: gut, Hp: hepatopancreas, O: ovotestis, W: whole snail, Hm: hemolymph, Ty: tray water, Tk: tank water.

We also investigated whether bacterial densities were linked between sample types, specifically whether higher hemolymph density would be associated with higher bacterial densities in the organs. However, we did not find consistent correlations between the bacterial densities in the organs, nor did we observe any correlation between hemolymph and organs bacterial densities (Supp. Table 3). This suggests that bacterial densities are likely regulated independently in adjacent organs and hemolymph.

## DISCUSSION

### 1. Snail organs and hemolymph have diverse and distinct microbiomes

Our analyses reveal the distinct and specific microbiomes associated with the stomach, gut, hepatopancreas, and ovotestis organs of *Biomphalaria sp.* snails. These organ microbiomes clustered separately from the hemolymph microbiome, exhibiting lower bacterial richness and stronger dominance. Microbial richness and dominance were primarily influenced by the sample type (organs, hemolymph or whole-snail), rather than by snail species. This suggests that organ habitat plays a strong role in shaping bacterial diversity. In addition, microbiomes tended to cluster according to organ functions, suggesting potential microbiome specializations. This pattern mirrors findings in transcriptomic analyses of different mammalian tissues, where gene expression clusters primarily by organs type rather than by species [29]. This may reveal interesting parallels between transcriptomic and microbiome fields, where units (expressed genes, microbes) could generate metabolic networks that define organ functions.

These specific organ and tissue microbiomes highlight the importance of investigating microbiomes at the organ level rather than the whole-organism level to obtain a more precise and robust understanding of microbiome compositions. While the investigation of organ microbiomes is common in vertebrates due to their size, this approach has received limited attention in small invertebrates. A few studies have explored organ microbiomes in mollusks (oysters, mussels [7–9]) and crustaceans (terrestrial crabs [10] and isopods [11]). Interestingly, for some - but not all - of these organisms, similar microbial patterns to those observed in snails were noted. For instance, hemolymph and gut microbiomes tend to cluster by sample types rather than by species in oysters (*Crassostrea gigas*) and Mediterranean mussels (*Mytilus galloprovincialis*) [7]. Additionally, the hemolymph microbiome showed higher observed richness than the gut, and this organ exhibited similar richness to that of the surrounding water. Another analysis of the Mediterranean mussel showed specific microbiomes for the hemolymph, gill, stomach, hepatopancreas and foot, all distinct from the water environment [8]. However, bacterial richness in the mussel hemolymph was similar to that in the stomach [8]. In crustaceans, the terrestrial brachuryan crab *Chiromantes haematocheir* showed specific organ microbiomes (in several gut sections, gills, and gonads) with a very limited bacterial diversity compared to the environment [10]. Similar to our findings, the organ microbiomes from different crab populations clustered by organ rather than by host population. The terrestrial isopod *Armadillidium vulgare* also showed a limited bacterial diversity in organs (nerve cord, digestive systems, gonads, and hemolymph) compared to environmental samples. However, their microbiomes clustered by population (field or laboratory) rather than by organs. While some of these differences may be attributed to technical factors (sample processing or analysis pipeline), they may also highlight species-specific strategies for managing organ microbiomes, which requires further investigation.

Snail microbiome studies have primarily focused on the gut (see [30] for a review). This organ is of particular importance, because its microbiome is involved in nutrient acquisition, which may have a significant impact on host and parasite fitness, and may ultimately influence species evolution [31]. We found that the snail stomach and gut have the highest bacterial densities per host cell, yet their microbiomes exhibited the lowest α-diversity indices. The high bacterial densities may be attributed to the larger available space for bacterial growth in the digestive tract. The significant difference in microbiome diversity (both α and β) between the digestive system and hemolymph provides an opportunity to investigate how microbiomes respond to pathogens. Snails could serve as an excellent model due to their susceptibility to a broad range of pathogens and parasites [32]. Furthermore, pathogens can inhabit various snail tissues. For example, *Angiostrongylus* nematodes develop in the gut, while schistosome trematode larvae migrate and develop in the hepatopancreas and ovotestis, both of which are bathed by the hemolymph. The variations in organ microbiomes and pathogen diversity provides an avenue to explore microbiome perturbation and adaptation both at site of infection (local) and at a systemic (distal) scale.

### 2. Whole snail: the importance of the shell

The whole-snail microbiome is another major area of study and was used for the first characterization of the *Biomphalaria* microbiome [33]. This approach is still in use today [34–40]. While sequencing a single “whole microbiome” per sample is cost-efficient and provides a snapshot of the diversity within the entire organism, it may also mask specific microbiome dynamics. In our study design, we included the analyses of whole-snail microbiomes to assess how representative they were compared to organ microbiomes. Whole-snail microbiomes exhibited one of the highest levels of α-diversity, comparable to that of the hemolymph microbiome. Interestingly, these whole-snail microbiomes were more similar in composition to the hemolymph microbiomes than to the composite of organ and hemolymph microbiomes. Hemolymph, therefore, could serve as a proxy for investigating whole-snail microbiome diversity, potentially enabling longitudinal studies of individual snail microbiomes through regular bleeding.

While studying whole-snail microbiomes can address specific questions, the results might be influenced by sample preparation methodology. Two strategies for preparing whole-snail samples are commonly used: either keeping the snail shell intact or removing it before homogenizing the snail bodies. This choice can significantly impact microbiome results, making direct comparisons between studies challenging. Shell removal is usually done by peeling the shell [33,35–37], which offers three main advantages: (i) it reduces potential contamination from the shell, (ii) facilitates sample homogenization (which can be challenging with hard shells), and (iii) limits PCR reaction inhibition by reducing the carryover of inhibitors such as calcium ions. However, shell peeling is likely to result in snail bleeding due to snail body retraction or tegumental damage. In the present study, we demonstrated that the hemolymph microbiome exhibits the highest α-diversity, and that the microbiome of whole snails (prepared with the shell intact) showed similar α-diversity to the hemolymph microbiome. Additionally, the whole-snail microbiome was more similar to the hemolymph microbiome than to organ microbiomes in terms of β-diversity, and a significant number of ASVs were shared between the whole-snail and hemolymph microbiomes. Hemolymph bleeding during shell removal could result in significant loss of bacteria, potentially limiting the capture of the full diversity of the whole-snail microbiome. Therefore, to better understand the impact of sample preparation on microbiome diversity (α, β and taxonomic), it is necessary to compare whole-snail microbiomes prepared with and without the shell.

### 3. Hemolymph and ovotestis: hotspots of microbial diversity?

Hemolymph and ovotestis harbored the highest number of total ASVs and distinct ASVs compared to the other snail organs (Fig. 5). This finding is unexpected, considering that the hemolymph contains immune cells responsible for targeting and eliminating microbes. While the precise role of the hemolymph microbiome, if any, remains unknown, the maintenance of a diverse microbiome through tolerance mechanisms may serve to control microbial populations by promoting competition between microbes and limiting immune reactions to prevent host damage, as observed in the gut of vertebrates [41]. The presence of a diverse microbiome in the ovotestis is also surprising due to remote anatomical position at the center of the coiled shell. This location should minimize its exposure to the hemolymph and other organs and their respective microbiomes. The ovotestis remote position could explain its lower bacterial density compared to other organs. However, this low bacterial density, coupled with high diversity, likely contributes to the ovotestis microbiome’s variable clustering pattern relative to other organ microbiomes (Fig. 3). It either clusters with the neighboring hepatopancreas in *B. alexandrina*, or separately from other organs in *B. glabrata*. While the exact role of the ovotestis microbiome is still undefined, we hypothesize that its diversity may be linked to providing nutrients and metabolites essential for egg production, such as vitamin E [42]. Understanding the drivers of this diversity could be crucial for developing strategies aimed at disrupting organ microbiomes as a means to control snail populations.

The ovotestis microbiome also exhibited a high proportion of unassigned taxa, a characteristic shared with the hepatopancreas and, to a lesser extent, the hemolymph. The proportion of unassigned taxa was notably higher in *B. alexandrina* compared to *B. glabrata*. Further characterization of these unidentified bacteria will be essential to better understand their potential contribution to ovotestis physiology.

#### 4. Implications of organ specific microbiomes

Snails are vectors of a variety of trematode parasites [32]. Over several weeks of host infection, these parasites undergo different developmental stages, migrating through snail tissues and developing within various snail organs. We previously hypothesized that parasites are exposed to the hemolymph microbiome [15]. However, our current findings indicate that parasites are exposed to an even more diverse range of microbiomes within snail organs. This observation raises three key questions: (i) How do these host microbiomes respond to the presence of parasites, and are there organ – or tissue - specific reactions? (ii) How do host microbiomes influence the parasites and their life-history traits (e.g. compatibility, development time, cercariae production, etc.)? (iii) Do parasites acquire microbiomes from the host? To date, only a few studies have investigated the impact of parasite infection on snail microbiome [37,40,43], with most focusing on whole-snail microbiomes or feces. However, this whole-organism approach may obscure specific changes occurring within the organ or hemolymph microbiomes, limiting our understanding of the microbiome’s role in parasitic interactions.

In addition of bacteria and archaea, the presence of microeukaryotes in *B. glabrata* has recently been investigated [38]. In this study, authors showed co-variation of microeukaryotes and bacteria in association with planorbid species. While this represents a significant advancement, as microeukaryotes is often overlooked, the observed co-variation was measured at the whole-snail level. These microeukaryotes could potentially exhibit organ-specific distributions, similar to those observed for bacteria. Investigating these distributions could provide future insights into this co-variation and reveal subtler patterns that have yet to be identified.

The presence of microbiomes associated with organs raises important questions about localization and colonization. While we can expect the microbes of the digestive system to be mainly localized in the lumen of the organs and those of the hemolymph to be free-floating, the localization of microbes in the hepatopancreas and ovotestis remains to be determined. While insects have specialized structure (bacteriocytes) to harbor symbiotic microorganisms [44], it is unknown whether snail microbes are randomly localized on the organ surface or if they tend to occupy specific areas within the organs, such as ducts or other dedicated structures. Additionally, the mode of microbial colonization of these organs is still unclear. We showed that some microbes are ubiquitous in snails and not detected in the environment, suggesting widespread dispersion within the host, possibly through connection between organs or leakage. Tracking labeled microbes could be a useful approach to investigate the pathways they use to colonize their host [45].

While some microbes are ubiquitously found across all organs and hemolymph, the majority of remaining microbes are distinct to each compartment. This suggests that microbiome homeostasis is maintained within each organ and hemolymph. This maintenance might be passive, with specialized microbes thriving in these different environments, or more active, involving specific mechanisms used by the snail to select them. Potential co-adaptation between snails and their microbiomes might be genetically mediated, as a snail genetic locus has been associated with the abundance of specific microbes using whole-snail microbiome analysis [35]. Conducting similar association studies at the organ or hemolymph level could be more powerful for detecting genetic determinants controlling host-microbe associations.

Distances between whole-snail microbiome communities of different snail species have been found to correlate with genetic distances between these species, suggesting phylosymbiosis [36]. Phylosymbiosis has been demonstrated in both invertebrates and vertebrates [46], and can also occur within different sections of the same organ in vertebrates (e.g. along the length of the gastrointestinal tract in rodents [47]). Similar observations could likely be made in invertebrates. The presence of diverse, organ-specific microbiomes in snails provides an excellent model for investigating phylosymbiosis at the organ level in invertebrates.

### 5. Limitations

Our characterization of snail organ microbiomes revealed diverse and organ-specific microbiomes. However, some of our findings may be influenced by aspects of our study design. Firstly, we did not control for snail age in both species, which may explain some of the observed variation in microbiomes. While we sampled snails of similar sizes, size correlates with age but is also influenced by population density [48]. As the dynamics of snail microbiomes during aging are unknown, it is possible that some of the differences observed between or within species are due to snail age and physiological state. For instance, the greater variation in bacterial density observed in the hepatopancreas of *B. glabrata* could reflect physiological differences, such as gut leakage.

Secondly, we sampled snail microbiomes at a single timepoint, which limited our ability to assess the stability of microbiome differences over time. Nevertheless, we think our results should be robust to short-term variations, as we integrated morning and afternoon sampling to mitigate potential effects of daily chronobiological variations, and sampled two independent tanks to minimize batch effects.

Lastly, all snails from the both species were raised in similar environmental conditions (i.e., exposed to the same water and food sources). This similarity in environmental conditions may partially explain why the organ, rather than species, is the primary factor responsible for microbiome differences. To confirm the preeminence of the organ factor in shaping snail microbiome diversity, we plan to analyze organ microbiomes from different snail species co-existing in diverse settings or natural locations, following the approach used in studies of bivalve mollusks [7] and crabs [10].

## CONCLUSIONS

We characterized the specific microbiomes associated with the hemolymph and organs (ovotestis, hepatopancreas, gut, and stomach) of two species of *Biomphalaria* snails, vectors of schistosome parasites. These microbiomes showed higher similarity based on the sample types rather than host species, suggesting that similar factors are shaping these microbiomes across different hosts. The hemolymph exhibited the highest microbial diversity, while the gut and stomach showed the highest bacterial load among the organs. Comparing these microbiomes to whole-snail microbiomes revealed that the majority of microbial diversity in whole-snails was shared with the hemolymph, though the whole-snail microbiome did not capture all the diversity present.

These findings will significantly impact investigations into the role of microbiomes in the interaction between snails and their parasites. The composite nature of the whole-snail microbiome, one of the most studied sample types in the field, likely obscures critical microbiome dynamics that occur at a more localized level. Our study advocates for a reevaluation of the whole-snail sample type and provides valuable insights to guide future experimental designs for more accurate and reliable microbiome studies.

## DECLARATIONS

### Ethics approval and consent to participate

Not applicable

### Consent for publication

Not applicable

### Availability of data and materials

Raw sequencing data are accessible from the NCBI Sequence Read Archive under BioProject accession number PRJNA1113672. Commands and scripts used for processing sequencing data and performing downstream analysis are available in a Jupyter notebook on Github (https://github.com/fdchevalier/Biomphalaria-organ-microbiomes).

### Authors’ contributions

LVC, TJCA, FDC, and WL designed the experiments. LVC prepared and collected the samples, extracted DNA and prepared the 16S libraries and sequencing run. SCN performed the qPCR assays. LVC and FDC performed all the data analyses. LVC, FDC and WL wrote the first version of the manuscript. All authors edited the manuscript and approved the final version.

## Acknowledgements

We thank Poppy Lamberton and Lisa Ranford-Cartwright for supporting LVC in the opportunity and funding acquisition for her internship to Texas Biomed Research Institute. We thank the Biomedical Research Institute (<u>NIAID Schistosomiasis Resource Center</u>, Rockville, MD) for providing the BgBS90 snails through NIH-NIAID Contract HHSN272201700014I, and the Theodor Bilharz Research Institute (Giza, Egypt) for providing *B. alexandrina* snails. We thank Dr. Shelley Cole and Vanessa Ayala (Texas Biomedical Research Institute) for their help with MiSeq sequencing.

## SUPPLEMENTARY FILES

**Supp. Fig. 1 – Rarefaction curves.** Curves were plotted for each sample type (Hm: hemolymph, S: stomach, G: gut, Hp: hepatopancreas, O: ovotestis, W: whole snail, Ty: tray water, Tk: tank water). Each of these curves plateau demonstrating that sampling more reads (x axis) would not have increased ASVs richness (y axis).

**Supp. Fig. 2 – α-diversity statistical results.** We performed Wilcoxon pairwise comparisons between the different sample types of *B. alexandrina* (Ba) and *B. glabrata* (BgBS90) for each α-diversity index (Fig. 2). The p-values are not adjusted for multiple comparisons. Results in these comparisons are similar to the linear mixed-effect model results (Supp. Table 3).

**Supp. Fig. 3 – Principal component analysis of snail hemolymph, organs, whole snails, and water.**

**Supp. Fig. 4 – Taxonomic diversity of sample types in each snail species.** Relative abundance of the different classes in each sample type for each snail species (A & B).

**Supp. Fig. 5 – Shared ASVs between sample types by species (full set).**

**Supp. Fig. 6 – Proportion of shared ASVs between sample types in pairwise comparisons.** The upper and lower triangles correspond to sample types of *B. alexandrina* (Ba) and *B. glabrata* (BgBS90), respectively.

**Supp. Fig. 7 – Statistical results 16S rRNA gene density.** We performed Wilcoxon pairwise comparisons between the organs/whole snails or hemolymph/water of *B. alexandrina* (Ba) and *B. glabrata* (BgBS90) for the 16S density (Fig. 7). The p-values are not adjusted for multiple comparisons. S: stomach, G: gut, Hp: hepatopancreas, O: ovotestis, W: whole snail, Hm: hemolymph, Ty: tray water, Tk: tank water.

**Supp. Table 1 – Earth Microbiome Project primers used for generating each library.**

**Supp. Table 2 – Library statistics.**

**Supp. Table 3 – Linear mixed-effect model results.**

